# Programing Immunogenicity of Dengue EDIII Vaccine Antigens Using Engineered Outer Membrane Vesicles (OMVs)

**DOI:** 10.1101/2025.02.25.640071

**Authors:** Srinivas Duvvada, Farhan Ahmed, Rafiq Ahmad Khan, Shaikh Matin Rahim, Saima Naaz, Aradhna Mariam Philips, David Putnam, Avery August, Nooruddin Khan

## Abstract

Dengue is a mosquito-borne viral infection and is more prevalent in the world with no therapeutics and suboptimal vaccine performance against all four serotypes of the dengue virus. Hence, there is an urgent requirement for a non-infectious and non-replicative vaccine candidate that can elicit a balanced and serotype-specific immune response. In this study, we have engineered bacterial outer membrane vesicles (rOMVs) that display EDIII antigens (EDIII rOMVs). The current formulation modulates the expression of costimulatory molecules on antigen-presenting cells (APCs) as well as enhances the uptake and presentation. Subsequently, the EDIII rOMVs elicited a strong antigen-specific polyfunctional response from CD4+ and CD8+ T cells. The robust antibody response was facilitated by a germinal center reaction characterized by high T follicular helper (Tfh) and B cell response levels in the mice that received EDIII rOMVs. Notably, the produced antibodies demonstrated the ability to neutralize all four dengue virus serotypes in an in vitro infection model, indicating its potential role in protective immunity.

**Graphical abstract:** 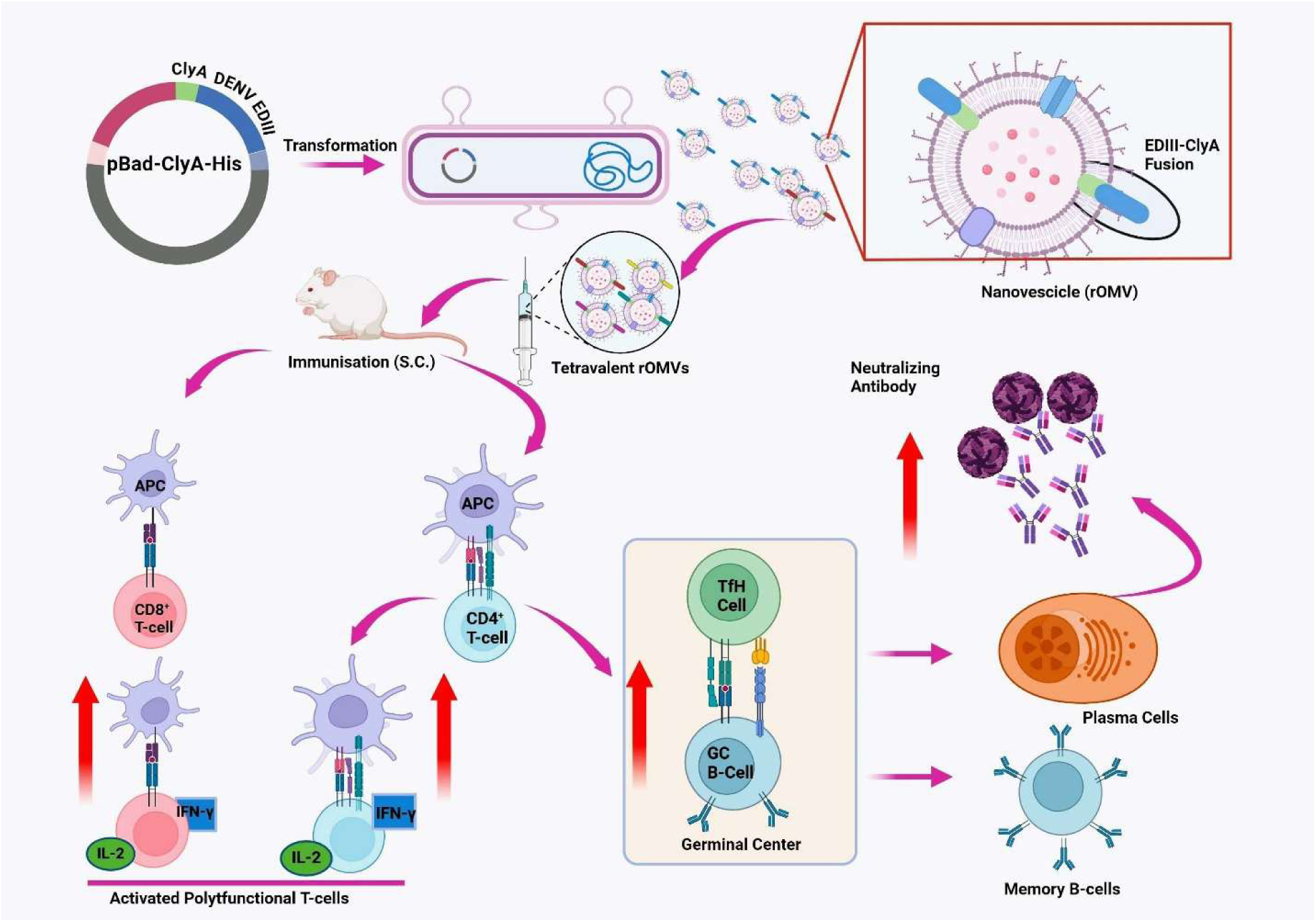

## Introduction

Dengue is transmitted by the infected mosquito *Aedes agypti* and caused by four closely related serotypes of dengue virus (DENV). DENV belongs to the family Flaviviridae and includes other clinically important viruses, such as Yellow Fever Virus (YFV), West Nile virus (WNV), Zika virus (ZIKV), and Japanese Encephalitis virus (JEV). In some cases, DENV can lead to life-threatening conditions, such as dengue shock syndrome (DSS) and dengue hemorrhagic fever (DHF). Dengue infection spreads at an alarming rate, and it is estimated that 390 million cases occur every year[1]. Infections are increasing, mainly due to the unavailability of antiviral therapeutics against DENV and the lack of a potent tetravalent vaccine. Dengue poses an enormous economic burden in addition to health problems[2]. Given the lack of effective antiviral therapies against DENV, a vaccine that protects against all four DENV serotypes is urgently required.

Development of vaccines against dengue is marked by considerable challenges, primarily due to various underlying factors, such as the presence of four distinct serotypes of the dengue virus (DENV 1-4), each capable of causing disease, and antibody-dependent enhancement (ADE) phenomenon, whereby pre-existing antibodies from a previous DENV infection can exacerbate the severity of subsequent infections of different serotypes. To address this, vaccine candidates must elicit neutralizing antibodies against all four serotypes, which considerably complicates the development process [3]. Despite these challenges, many vaccines for dengue are in preclinical and clinical development stages, and one vaccine is already licensed in many countries. The vaccine developed by Sanofi Pasteur (brand name Dengvaxia (CYD-TDV)) is a live attenuated chimeric vaccine with a 17D backbone, in which the prM and E proteins of DENV serotypes (1-4) have been exchanged for the proteins of YF17D[4]. Dengvaxia is effective only in people who are exposed to dengue previously and is not recommended for use in seronegative individuals, as it has been observed that vaccination increases the risk of severe infection in individuals who do not have preexisting immunity to dengue[5]. In addition to Dengvaxia, many live attenuated vaccine candidates have shown promising results in clinical trials; however, data on their ability to provide long-term protection vaccines are not available[5].

Although live attenuated vaccines provide long-lasting memory responses, there is always a risk of them regaining their virulence. One of the major challenges faced by live attenuated vaccine developers in the case of dengue is obtaining a balanced replication of components of all DENV serotypes[6]. Furthermore, these vaccines generate antibodies that lead to ADE[7]. Alternative approaches include developing non-replicating subunit vaccines that are safe and relatively easier to generate vaccine components for all serotypes in a balanced manner. DENV Envelope protein domain III (EDIII) has been considered a potential subunit vaccine candidate for dengue owing to its ability to produce neutralizing antibodies[8], potentiate immunogenicity, require boosters and adjuvants[9],and susceptibility to degradation by host proteases. Therefore, an effective delivery and adjuvant system is required to overcome these limitations. Nano-based delivery systems offer enormous advantages and are used to enhance the immunogenicity of subunit vaccines. Such nanocarriers enhance the immunogenicity of vaccine antigens by preventing degradation, increasing uptake, elicit sustained release, and target immunogens to Antigen Presenting Cells (APC)[10, 11].

Here, we used an outer membrane vesicle (OMV) based vaccine platform to enhance the immunogenicity of the DENV EDIII antigen. OMVs are released by gram-negative bacteria during growth both in vitro and in vivo[12]. These bacterial nanoparticles are bilayered and spherical, containing many components on their external surfaces [13, 14]. OMVs have a wide variety of bacterial antigens on their surface[15], and they have an ideal size that facilitates their uptake by immune cells. In addition, OMVs have immunopotentiators such as TLR agonists, which bring about the self-adjuvanticity of OMVs. This property of OMVs makes them strong activators of innate immune response[16, 17]. OMV-based vaccines are approved for use in humans, most notably the formulation approved in the European Union and by the FDA[18] (Bexsero®, Novartis) to prevent Neisseria meningitidis serogroup B infection. In the current study, we used recombinant OMVs (rOMVs) in which the protein of interest (EDIII) was fused to a membrane protein anchor, which then co-localized this fused protein to the outer membrane to generate rOMV displaying EDIII. We expressed EDIII of four serotypes of DENV separately in rOMV and physically mixed these vesicles to develop a tetravalent formulation. The immunogenicity of this formulation was tested in BALB/c mice, and we observed strong antigen-specific T and B-cell responses along with the neutralizing capability of antibodies in an in vitro infection model.

## Results

### Successful design and characterization of ClyA-EDIII recombinant OMVs

Recombinant OMVs are made based on a simple technique in which the protein of interest is fused to a membrane protein anchor, which then co-localizes the fused protein to the outer membrane of vesicles. We used one such example of a transmembrane protein, ClyA, a pore-forming cytotoxin in the outer membrane of OMVs of enterobacteria such as *E. Coli*, particularly the ClearColi® strain with diminished endotoxin content, to mitigate adverse immune reactions. We generated fusion proteins between candidate antigens and Cly A[23, 24], expressing EDIII antigens fused with ClyA from DENV serotypes, individually and physically mixed to form a tetravalent formulation. We first cloned DENV EDIII into the pBAD-ClyA vector to produce rOMVs (Fig. S1). The EDIII gene was cloned at the C-terminal of the ClyA protein to display the antigens on the surface of OMVs, as previously performed with different antigens[25]. Expression of these antigens on purified rOVMs was confirmed by western blotting using antisera against all four EDIII antigens (Fig. S2). Isolated rOMVs displaying DENV EDIII of all four serotypes were characterized for their shape and size by TEM, and the results showed that vesicles were spherical and less than 100 nm in size (Fig. 1C). The size distribution of vesicles was analyzed by DLS, and data have shown that the average size of rOMVs is less than 100 nm, with polydispersity in nature. (Fig. 1C lower panel).

**Figure 1.**
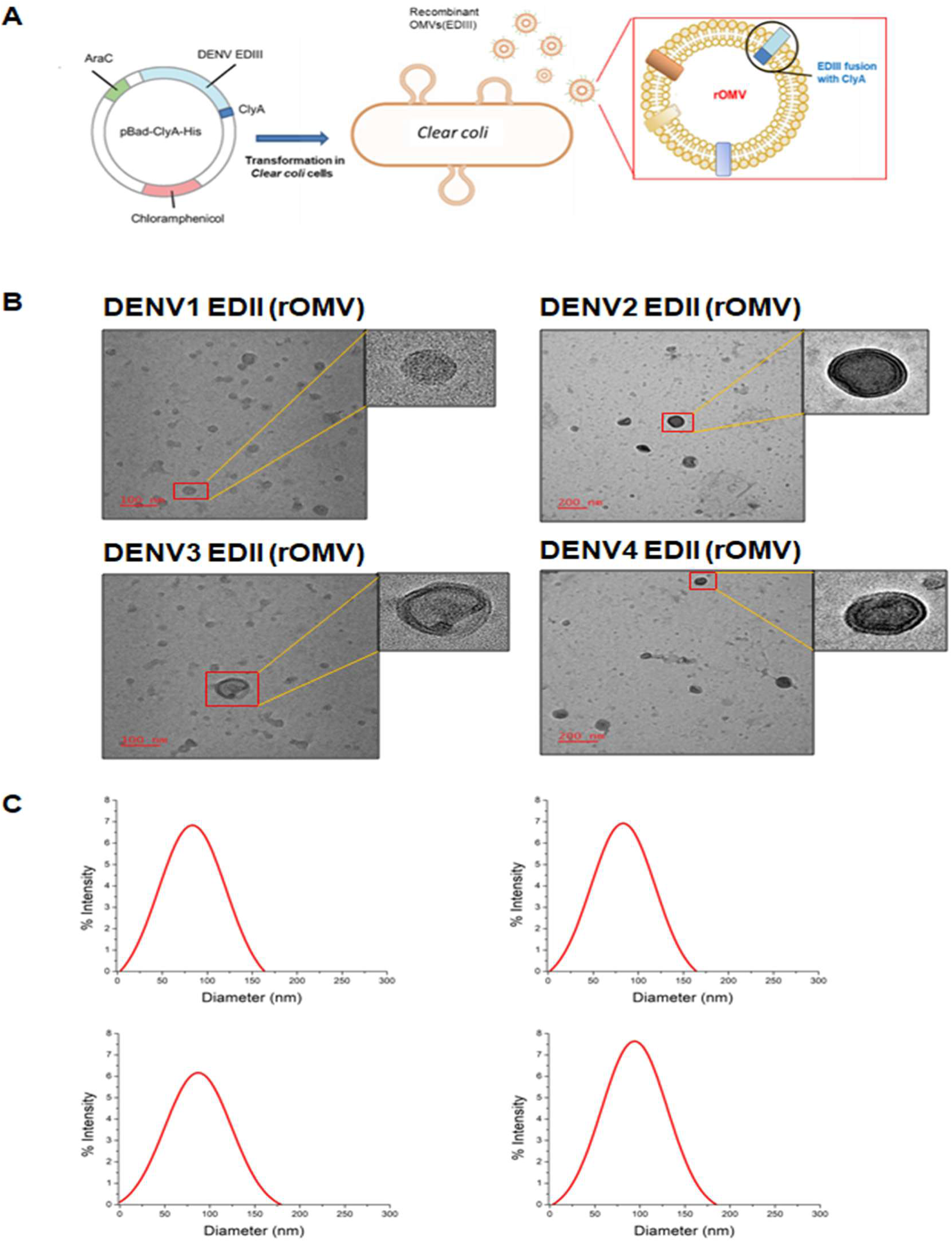
Expression, purification and characterization of EDIII containing rOMVs. A) EDIII antigen was cloned into pBAD-ClyA-His Vector and transformed into ClearColi cells, and OMVs with surface displayed antigen isolated. B) & C) Purified rOMVs containing all the four antigens were characterized for shape and size using Transmission Electron Microscopy (TEM) and Dynamic light scattering method (DLS) below.

### rOMVs augmented the antigen uptake, presentation and expression of co-stimulatory molecules on Antigen Presenting Cells an in-vitro system

The innate effector functions of antigen-presenting cells (APCs), including antigen uptake, processing, presentation, and co-stimulation, are crucial for programming effective and durable antigen-specific adaptive immune responses [26, 27]. To assess the potential of rOMVs to modulate these key signals, we investigated their ability to enhance antigen uptake, presentation, and expression of co-stimulatory molecules on antigen-presenting cells (APCs). Therefore, Dendritic cells (DCs) were incubated with GFP-expressing rOMVs at two different temperatures (37°C and 4°C) and analyzed using flow cytometry (FACS). The results demonstrated the efficient uptake of rOMVs at 37°C, whereas no uptake was observed at 4°C, suggesting the involvement of active endocytosis[28] in rOMV internalization (Fig. 2A Fig. S4). Following antigen uptake, presentation of antigens on major histocompatibility complex (MHC) molecules is a critical step in T-cell priming. We evaluated the expression of MHC class I (MHCI) and MHC class II (MHC II) molecules in DCs after stimulation with rOMVs. Significant upregulation of both MHCI and MHC II molecules was observed (Fig. 2B; Fig. S3), indicating that rOMVs could effectively prime DCs for antigen presentation. Co-stimulatory molecules, such as CD80 and CD86, are essential for the activation of naïve T-cells and the establishment of an effective adaptive immune response[29]. Stimulation of DCs with rOMVs resulted in the enhanced expression of both CD80 and CD86 (Fig. 2B; Fig. S3). This upregulation underscores the capacity of rOMVs to activate DCs and promote T cell co-stimulation. These findings established that rOMVs effectively modulate critical aspects of innate immunity, including antigen uptake, antigen presentation, and co-stimulatory signaling.

**Figure 2.**
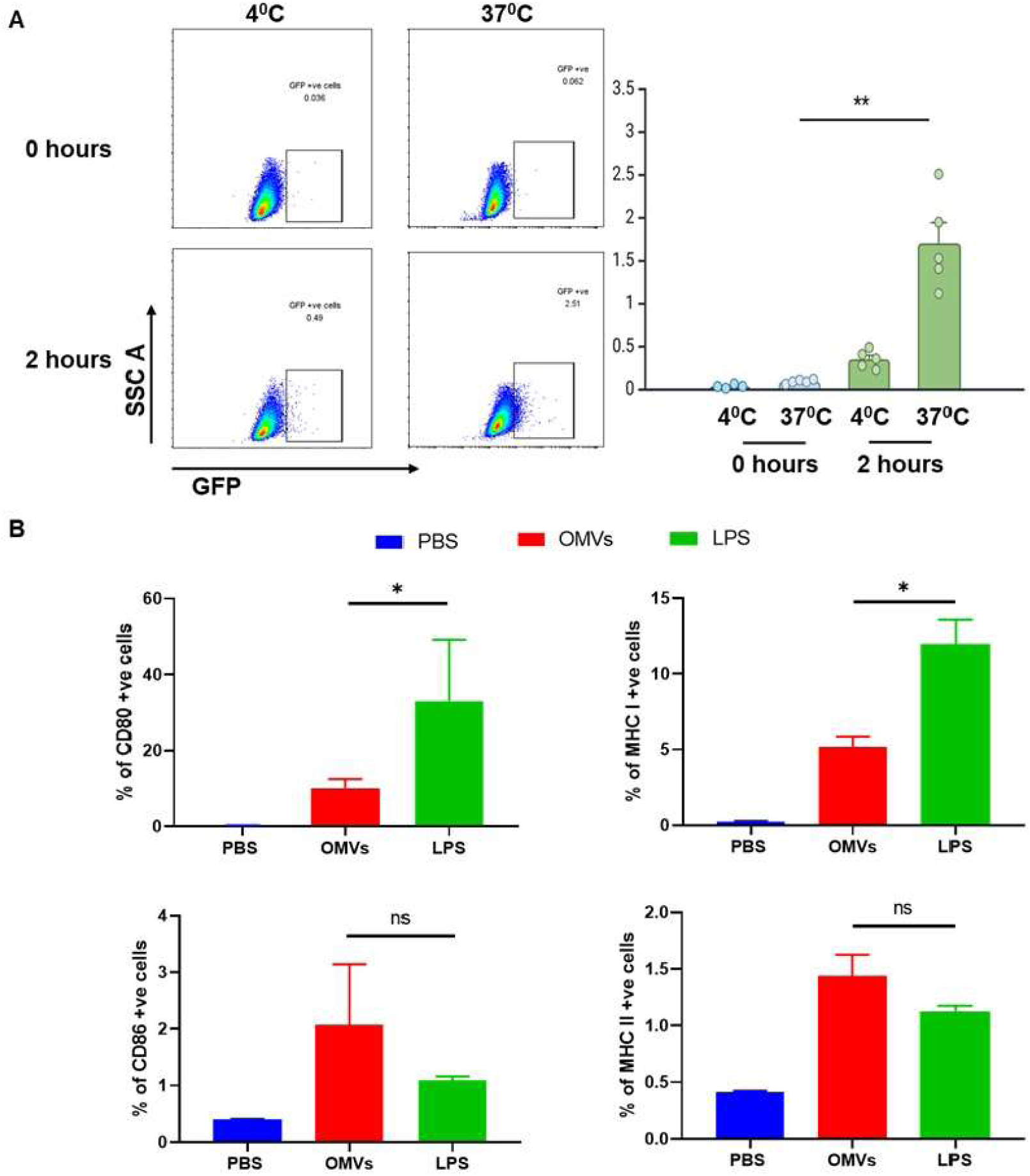
Assessment of antigen uptake and co-stimulatory molecules expression induced by rOMVs. (A) Antigen-presenting cells (DCs) were incubated with GFP-expressing rOMVs at two different temperatures (37°C and 4°C). The percentage of GFP-positive cells was evaluated using flow cytometry (FACS), demonstrating significant rOMV uptake at 37°C while no uptake was observed at 4°C. (B) The expression levels of co-stimulatory markers (CD80 and CD86) and antigen presenting molecules (MHCI and II) on DCs post-rOMV stimulation and FACS analysis.

### rOMVs expressing DENV EDIII induce robust antigen-specific T cell response in the mice

After successfully designing and characterizing rOMVs displaying EDIII, we further explored the ability of rOMVs to stimulate antigen-specific polyfunctional T cell responses to the dengue antigen. Mice immunized with the different formulations were sacrificed after 28 days, and splenocytes were collected from the spleen and restimulated with the same antigen in the culture medium. Recall assay data showed that the rOMV(tEDIII) formulation induced a considerable degree of antigen-specific CD4+ and CD8+ T cell responses as compared to EDIII antigen alone and other control groups. Specifically, the frequency of polyfunctional T cells, which produce IL-2 and IFN-γ among CD4+ (Fig. 3 B-E) and CD8+ T cells (Fig. 3 G-J), was enhanced in animals immunized with the rOMV(tEDIII) formulation compared to free antigens or control groups. Polyfunctional T-cells are important for inducing effective and durable immunity against viral antigens.

**Figure 3.**
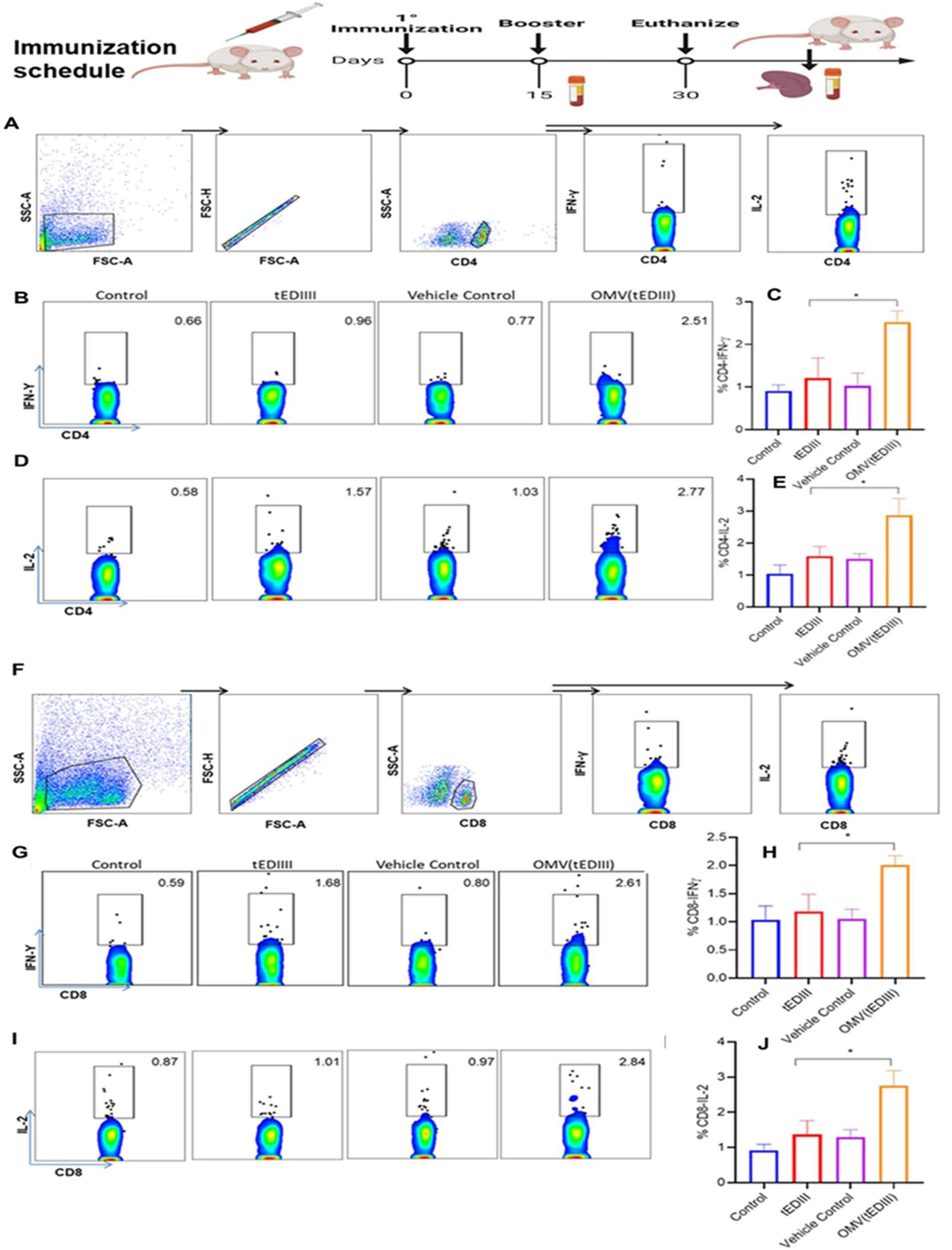
The tetravalent formulation of rOMVs expressing DENV EDIII from all the serotypes induces antigen specific CD4+ and CD8+T cell response in the mice. Splenocytes from immunized mice were stimulated with purified EDIII antigens and analyzed by flow cytometry. A and F) Gating strategy to distinguish cells population. B) and C, G) representative flow cytometry plot of frequency of IFN-γ secreting CD4+ cells; while G, H) and represents CD8+ Cells. Box plot (C) and (H) represent the average percentage of IFN-γ producing CD4+ and CD8+ cells respectively. D, and J) are representative flow cytometry plot of frequency of IL-2 secreting CD4+ and CD8+ cells. Box plot (E) and (J) represent average percentage of IL-2 producing CD4+ and CD8+ cells respectively in the various immunization groups; Control (PBS), tEDIII (soluble recombinant EDIII), Vehicle Control (empty OMVs), OMV (tEDIII)-OMV expressing EDIII from all the four DENV serotypes. Each data set is a representative of 10 animals (5×2) and data is presented as a mean with SEM. Value of significance *P < 0.05 is considered as significant data.

### rOMV expressing DENV EDIII in tetravalent formulation induces a significant antigen-specific antibody response

The effectiveness of a successful viral vaccine is measured by the propensity of the B cell response to produce an antigen-specific high-affinity antibody that ultimately helps neutralize the viruses[30]. Therefore, DENV EDIII-specific total IgG from serum was measured using indirect ELISA. The results showed that antigen-expressed rOMVs induced significantly higher antibody levels against all four distinct serotype antigens compared to the antigens alone or control groups after 14 days of priming. After the booster dose, the level of antibody was further enhanced on day 28 (almost twice the value of the fourteenth day) in the groups of animals that were immunized with rOMV (tEDIII) as compared to (tEDIII) alone or in control groups (Fig. 4A-D). Notably, OMVs in tetravalent combinations induce a type-specific balanced antibody response against all four serotypes, which is an encouraging observation for formulating an effective vaccine against dengue virus.

**Figure 4.**
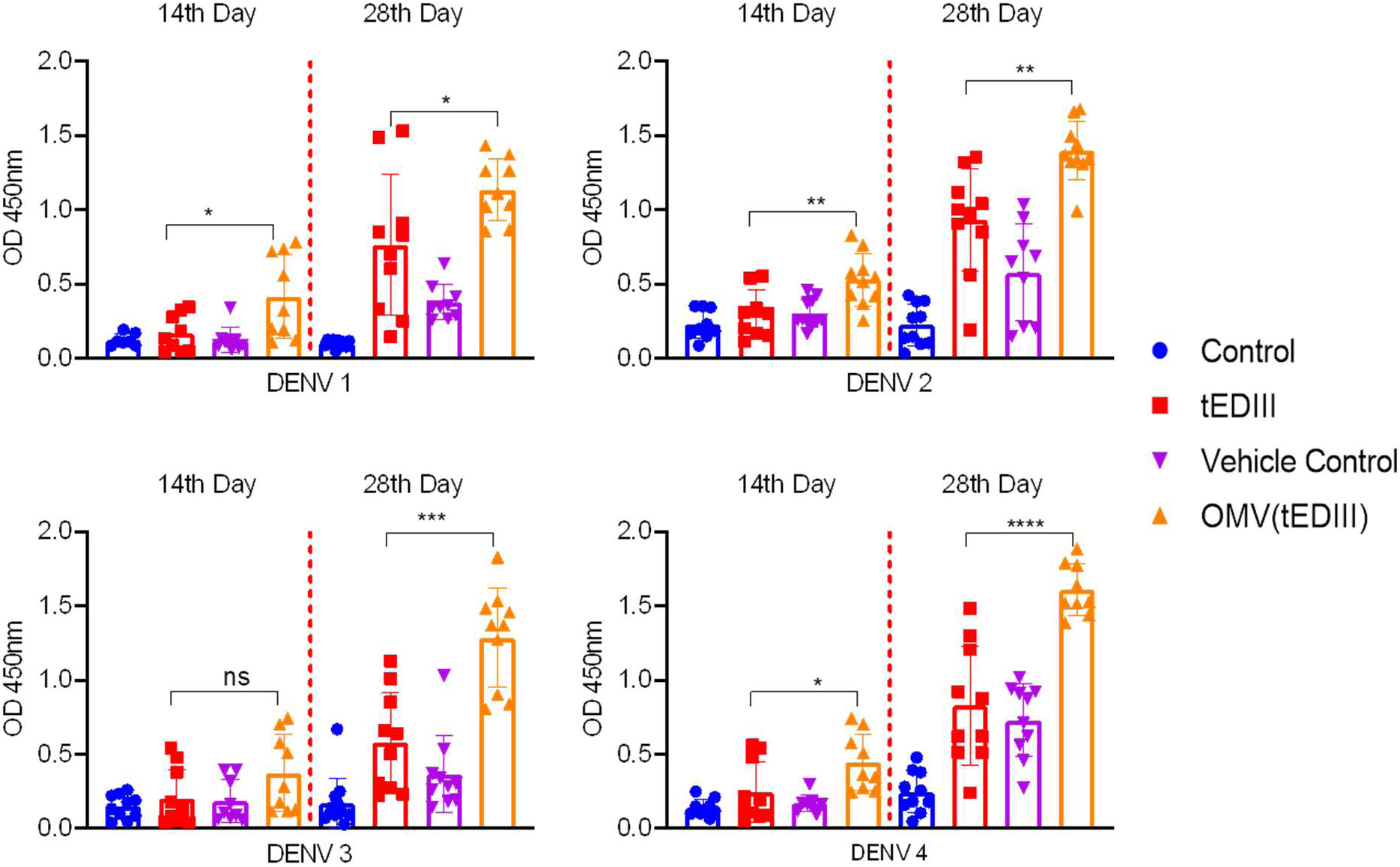
rOMV expressing DENV EDIII from all the four serotypes of DENV virus in a tetravalent formulation induces significant antigen-specific antibody response. Antigen-specific total IgG from serum of immunized mice was determined by indirect ELISA at day 14 and 28 post immunization. **A)** Antibodies against DENV1, **B)** Antibodies against DENV2, **C)** Antibodies against DENV3 and **D)** Antibodies against DENV4. Data of each groups contains 10 animal (5×2) and data represented as mean with SEM. Value of significance \**P* < 0.05, \*\**P* < 0.005, \*\*\**P* < 0.0005 and \*\*\*\**P* < 0.00005).

### Tetravalent (DENV 1-4) EDIII expressed in OMVs augmented Tfh and GC-B cell response

Strong humoral immunity plays a vital role in tackling viral infections by producing high levels of high-affinity antibodies. GCs are important transitory structures in the lymph nodes where specialized B cells differentiate into memory B cells, and long-lived antibody-secreting plasma cells with the help of Tfh CD4^+^ T cells[26, 31-33]. Therefore, in the current study, cells were isolated from the draining lymph nodes of immunized mice and stained with fluorophore-conjugated marker antibodies for Tfh CD4^+^ T cells (PD-1^+^, CXCR5^+^ and CD4^+^) and GC B cells (B220^+^, GL7^+^ and Ig^+^), followed by flow cytometric analysis. The results showed that the frequency of Tfh cells (PD-1^+^, CXCR5^+^, and CD4^+^) in the animals immunized with rOMV(tEDIII) formulation was significantly enhanced compared to that in the (tEDIII) or control groups (Fig. 5A-B). Tfh cells assist in B cell affinity maturation and differentiation in the germinal center. Therefore, we also measured the occurrence of GC B cells in the lymph node, and the results indicated that the abundance of GC B cells (GL7^+^ and Ig^+^) was higher in the rOMV(tEDIII)-immunized group than in the other groups (Fig. 5 C-D).

**Figure 5.**
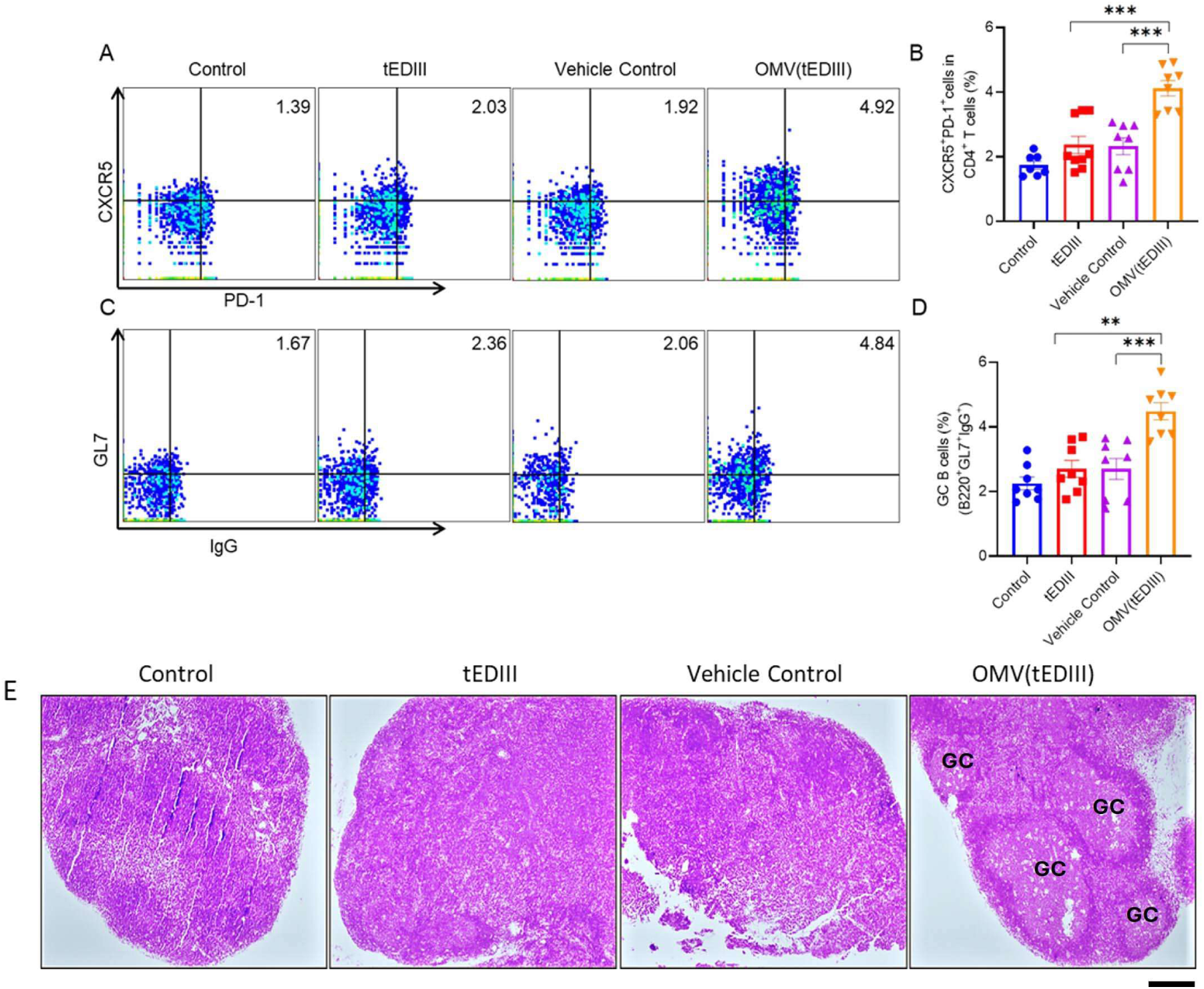
The tetravalent formulation of rOMVs expressing DENV EDIII from all the serotypes enhanced Tfh and GC-B cell response in the lymph nodes of immunized mice. Cells from inguinal lymph node were isolated and stained with Tfh and GC-B cell surface marker antibodies and analyzed in flow cytometer. A and B) The frequency of TfH cells were identified by triple positive T cells shown as FACS representative plot (CD4^+^, CXCR5^+^ and PD-1^+^ cells) and precent frequency. C and D) The frequency GC B cells represented as triple positive cells in FACS representative plot (B220^+^, GL7^+^ and IgG^+^) and precent frequency. Each data set contains 8 animals (4×2) and data represented as mean with SEM and significance **P<0.005, ***P<0.0005 was considered as significant, analyzed by Student t test followed by Mann-Whitney test. E) Histopathological analysis of Inguinal lymph node by H&E staining to determine the formation of germinal center (GC). bar size =100µm. Representative data of 8 animals (4×2).

### Serum from mice immunized with EDIII antigens expressing OMV (tEDIII) effectively neutralizes all the serotypes of Dengue

The neutralizing antibody (nAb) titer plays a crucial role in assessing the effectiveness of vaccine formulations against dengue and other viruses, and measuring the ability of nAbs to neutralize the virus is an important functional parameter for evaluating the potency of the candidate vaccine [34]. In line with these findings, we assessed the virus-neutralizing capability of serum antibodies obtained following immunization in mice using an *in-vitro* infection model. Active viruses of all four serotypes were mixed with the serum of fully immunized animals from all immunization groups in separate tubes and then added to the monolayer of Vero cells for infection, and the infected Vero cells were analyzed by flow cytometry. The results indicate that serum from the OMVs (tEDIII) group of mice was able to effectively neutralize all the serotypes of Dengue Virus by significantly inhibiting infectivity in Vero cells. Therefore, the data, expressed in terms of percent normalized infection and flow cytometry-based neutralization tests-50 (FNT50), suggest that the OMV (tEDIII) vaccine formulation could provide protection to the Vero cells compared to tEDIII alone or control groups (Fig 6. A-D). The significant potency of the nAbs from the OMVs (tEDIII)-immunized mice substantiates the fact that antigens displayed on rOMVs induce polyfunctional T cells that modulate a potent B cell response, thereby inducing high-titer nAbs production, which is effective against all serotypes of the dengue virus.

**Figure 6.**
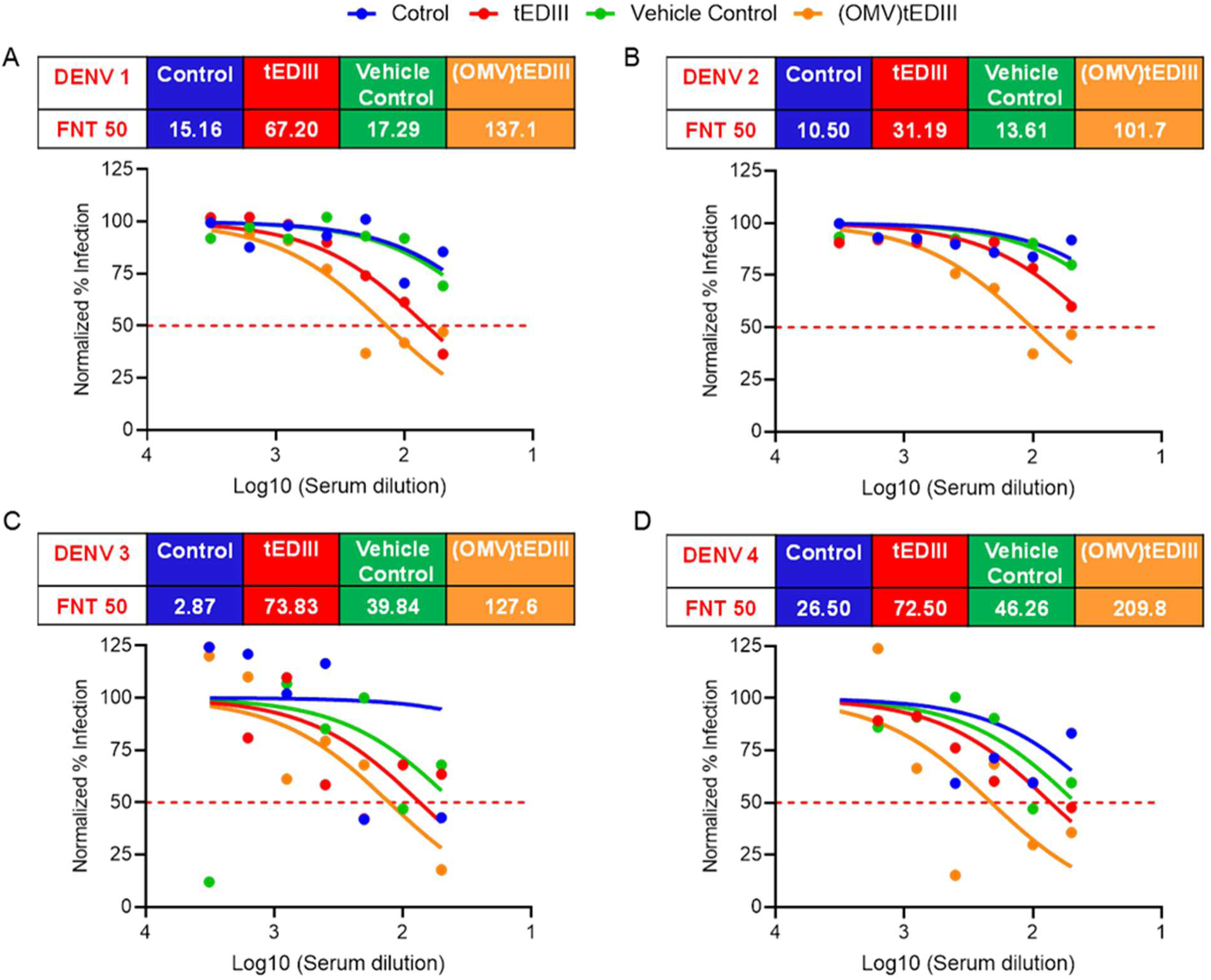
Serum from mice immunized with tetravalent formulation of rOMVs expressing DENV EDIII from all the serotypes effectively neutralizes all the serotypes of Dengue. The potential of immunized serum to neutralize dengue virus in vitro was assessed at various dilutions using the FACS neutralization test (FNT) for all four serotypes: DENV 1 (A), DENV 2 (B), DENV 3 (C), and DENV 4 (D). The FNT50 value was determined for each immunized serum sample, which indicates the serum dilution required to neutralize 50% of the virus, as determined by the FACS neutralization test for DENV 1 (A), DENV 2 (B), DENV 3 (C), and DENV 4 (D). The data is presented as the mean of multiple biological and experimental replicates, along with the standard deviation, and a significance level of p < 0.05 was considered significant, analyzed by student t-test followed by Mann-Whitney test.

## Discussion

The development of effective dengue vaccines remains a significant challenge for global health because of the complex nature of dengue virus (DENV) pathogenesis, including the existence of four serotypes (DENV-1 to DENV-4) and the risk of antibody-dependent enhancement (ADE). The limitations of the current dengue vaccines, including Dengvaxia®, highlight the need for alternative strategies. Although Dengvaxia is effective in seropositive individuals, it poses a risk of severe disease in seronegative recipients. Additionally, live attenuated vaccines face challenges related to balanced replication of serotypes and the potential for reversion to virulence. In this context, the current study presents a novel vaccine strategy using recombinant outer membrane vesicles (rOMVs) displaying envelope protein domain III (EDIII) of all four DENV serotypes. This tetravalent rOMV formulation demonstrated promising immunogenicity, addressing several challenges associated with dengue vaccine development.

The use of bacterial outer membrane vesicles (OMVs) as delivery platforms for antigens is a promising approach for vaccine development. OMVs naturally possess immunostimulatory molecules, including TLR agonists, which can activate antigen-presenting cells (APCs) and enhance both innate and adaptive immunity. The inclusion of pore-forming ClyA protein as a membrane anchor in this study ensured the efficient display of DENV EDIII antigens on the OMV surface. Transmission electron microscopy (TEM) and dynamic light scattering (DLS) analyses confirmed that the rOMVs maintained a spherical morphology with an average size below 100 nm, which is consistent with previous reports on their optimal parameters for cellular uptake and immunogenicity[33].

The ability of rOMVs to augment antigen uptake and presentation by dendritic cells (DCs) underscores their potential as efficient vaccine delivery systems. Flow cytometry analysis revealed that rOMVs were internalized through active endocytosis and significantly upregulated the expression of MHC class I (MHCI) and MHC class II (MHCII) molecules in DCs. Moreover, rOMVs increased the expression of the co-stimulatory molecules CD80 and CD86, which are critical for T cell activation. This modulation is pivotal for robust T cell priming and activation, consistent with previous reports highlighting the role of bacterial OMVs in bridging innate and adaptive immunity[35]. The ability of rOMVs to activate these innate immune attributes underscores their potential as versatile immunomodulatory agents for vaccine platforms.

A key feature of effective viral vaccines is their ability to elicit robust and durable T cell immunity[36]. The rOMV(tEDIII) formulation induced strong antigen-specific polyfunctional T-cell responses in immunized mice. Splenocyte recall assays demonstrated enhanced frequencies of CD4+ and CD8+ T cells producing IL-2 and IFN-γ, respectively. These cytokines are crucial for orchestrating cellular immunity, and are indicative of a potent and balanced immune response. Notably, polyfunctional T cells are associated with effective immunity against complex viral pathogens, including flaviviruses, and their presence suggesting the potential of rOMVs to generate long-lasting protective immunity. OMVs from different bacteria can activate innate effector responses [37, 38], which, in turn, provide a conducive signal for the effective priming of T cells that later define the outcome of immune responses.

The humoral response elicited by the rOMV(tEDIII) formulation was characterized by high levels of DENV EDIII-specific IgG antibodies against all four serotypes. Importantly, antibody responses were balanced across the serotypes, addressing a critical challenge in dengue vaccine development[39]. After a booster dose, antibody titers increased significantly, indicating the ability of the formulation to generate a strong anamnestic response. The balanced antibody response is particularly encouraging, as it may reduce the risk of ADE, a phenomenon that can exacerbate disease severity in subsequent DENV infections[40].

The ability of rOMVs to enhance T follicular helper (Tfh) cell responses and germinal center (GC) reactions further supports their potential as effective vaccine platforms. Tfh cells are essential for driving B cell affinity maturation and differentiation into plasma cells and memory B cells within germinal centers. This study demonstrated a significant increase in the frequency of Tfh cells in the lymph nodes of immunized mice. These findings indicate that rOMV(tEDIII) formulation can effectively promote the generation of high-affinity antibodies, which are crucial for neutralizing DENV and providing long-term immunity. Germinal centers have been linked to durable humoral immunity against various pathogen[41].

The most critical metric for evaluating the efficacy of a dengue vaccine is its ability to generate neutralizing antibodies (nAbs) that effectively inhibit all DENV serotypes. Serum from mice immunized with the rOMV(tEDIII) formulation demonstrated potent neutralization activity in vitro, as evidenced by the reduced infectivity of Vero cells exposed to live DENV of all four serotypes. Flow cytometry-based neutralization test (FNT50) results confirmed the broad-spectrum neutralizing capability of the induced antibodies. This underscores the potential of rOMV(tEDIII) as a tetravalent vaccine candidate, capable of providing protection against all DENV serotypes.

In contrast to current vaccine modalities, it possesses superior features such as non-replicating rOMVs, which eliminate the risk of reversion to virulence, making them inherently safer than live attenuated vaccines. The presence of natural immunopotentiators in OMVs reduces the requirement for additional adjuvants. The rOMV(tEDIII) formulation elicited a serotype-specific balanced antibody response, addressing a critical limitation of the existing vaccines. The simple production method for rOMVs allows cost-effective and scalable vaccine manufacturing. Although the current study provides compelling evidence for the immunogenicity of the rOMV(tEDIII) vaccine platform, several aspects warrant further investigation. First, the durability of the immune response and long-term protective efficacy must be evaluated in additional preclinical models and human clinical trials. Second, the potential for ADE should be rigorously assessed in future studies to ensure vaccine safety. Additionally, the scalability of rOMV production and the feasibility of combining this approach with other antigen delivery systems or adjuvants should be explored. The incorporation of novel adjuvants or immunomodulators can further enhance the immunogenicity and breadth of vaccine-induced immune responses.

## Conclusion

In conclusion, this study provides a new strategy for enhancing the immunogenicity of dengue vaccine antigens by using an rOMV-based self-antigen delivery and adjuvant system. Employing such approaches, we have developed a novel tetravalent nano-vaccine formulation comprising an immune-dominant EDIII antigen from all serotypes of DENV, which can confer strong vaccine-induced protective immunity capable of effectively neutralizing all serotypes of DENV (1-4). These findings highlight the importance of rOMV-based vaccine development platforms, which could be extrapolated to the development of effective, safe, and economical vaccine formulations against other emerging and re-emerging diseases.

## Materials and Methods

### Cloning, expression, and purification of rOMVs expressing EDIII from all the four serotypes of DENV

The cloned constructs for DENV EDIIIs in the pET28a vector were used as templates for cloning serotype-specific EDIIIs in the pBAD-ClyA vector to produce rOMVs. The restriction sites XbaI and HindIII (Fermentas, Thermo Fisher,USA) were introduced at the 5’ and 3’ ends of EDIII using forward and reverse primers. PCR was run for 35 cycles of denaturation (30 s, 95°C), annealing (30 s), and extension (25 s, 72°C) in a thermocycler (Applied Biosystems). Correctly sized amplification products were obtained for all four serotypes. The inserts obtained from the PCR were double-digested using XbaI and HindIII restriction enzymes. The pBAD-ClyA vector was also digested simultaneously using the same pair of restriction digestion enzymes to generate sticky ends compatible with the insert, and double-digested fragments were retrieved from the gel using Nucleospin gel/PCR cleanup (Machery Nagel). The digested insert was ligated to the vector using DNA Ligase (NEB), followed by transformation in E. coli DH5-alpha cells. Colonies were selected based on antibiotic resistance, and plasmids were isolated from selected colonies. Double digestion with the same pair of enzymes (XbaI and HindIII) was performed to confirm the clones. The digested plasmids were subjected to agarose gel electrophoresis, and bands corresponding to the size of EDIII were observed for all four DENV, types which confirm the successful cloning of EDIIIs in the pBAD-ClyA vector. After cloning, we expressed and purified rOMVs expressing EDIIIs according to previously described methods with modifications[19]. Briefly, the clones obtained were transformed into hyper vesiculated clear coli cells by heat shock. Single colonies were picked and inoculated into a liquid culture (LB broth) containing chloramphenicol (35 µg/ml) and grown at 37°C overnight in a shaking incubator. next day, 500 ml flasks containing LB broth and chloramphenicol (Himedia) were inoculated with overnight culture and kept in shaking incubator until OD600 reached ∼0.5. The expression was induced with L-arabinose (0.2%), and culture was allowed to grow for 16 h followed by centrifugation. The supernatants were then filtered through polyethersulfone membranes with a 0.22 μm pore size to remove any debris or cells. Subsequently, the supernatants were concentrated by ultrafiltration with a 100 kDa molecular weight cut-off (Vivaspin 50R, Sartorius). The retentate was rinsed once with 500 mL of PBS (pH 7.4), concentrated to 1 mL, and further subjected to ultracentrifugation at 150,000 × g for 90 min. Finally, the supernatant was discarded and resuspended in 500 ml filtered 1xPBS. The protein content of the final OMV suspensions was determined using the Bio-Rad Protein AssayTotal protein from rOMVs was determined by BCA estimation and an equal amount of protein (vesicles) was loaded in SDS-gel. rOMVs expressing EDIIIs were confirmed using anti-sera from EDIII-immunized mice, and anti-His was used to confirm empty OMVs.

### Physical characterization of DENV EDIII OMV

rOMV expressing EDIII were characterized by transmission electron microscopy (TEM) (TEM-FEI Tecnai) and Dynamic Light scattering (DLS) using (Anton Paar Pvt.ltd.). For TEM, the samples were suspended in water, contrasted with 2% uranyl acetate for two minutes, dried, and placed on a 200-mesh type-B carbon coated copper grid. The samples were observed under a TEM, and the data were collected and analyzed. DLS was used to determine the size distribution of the nanoparticles and OMVs in the suspension phase. The total protein concentration of OMVs was determined by bicinchoninic acid assay (BCA).

### Antigen Uptake and co-stimulation assay of rOMVs expression GFP and rOMVs

rOMVs expressing GFP were isolated from the media through a differential centrifugation process and incubated with Dendritic cells (1 × 10^5^ cells) seeded in 24 well plates and were performed at two different temperatures (4°C & 37°C) to check the phagocytic activity of DCs. After incubation for 2 h, media from the wells were removed and washed with PBS, and the cells were trypsinized and neutralized by adding RPMI (Gibco). DCs were collected from the 24 well plates into individual 1.5 ML tubes for further analysis. GFP-OMV-positive DCs were analyzed using a Flow Cytometer (BD Accuri 6^+^).

The expression of costimulatory molecules on antigen-presenting cells, such as dendritic cells, is key for immune modulation and the activation of adaptive immunity. To Check whether rOMV-treated DCs *in-vitro* can stimulate the expression of costimulatory markers, we treated DCs with LPS as a positive control and rOMVs and incubated them for 32 h and then stained with Anti CD80, CD86, MHC I, and MHC II fluorophore-tagged antibodies PE, PerCP, Alexa Fluor 647, and Alexa Fluor 488, respectively, added (1:250), and incubated for 1 h at RT. Finally, cells were washed and resuspended in FACS buffer, data were acquired using flow cytometry (BD Accuri C6), and data were further analyzed in Flowjo (BD bioscience).

### Animal experiments

All animal experiments were conducted in accordance with the animal ethics guidelines (Institutional Animal Ethics Committee/CPC SEA) of the University of Hyderabad. Six-to-eight-week-old Balb/c mice of the same age were immunized subcutaneously at the base of the tail in the following groups: (i) PBS control; (ii) soluble recombinant purified antigens (tEDIII) from all four DENV serotypes; (iii) empty OMVs (Vehicle Control); (iv) tetravalent formulation of OMVs expressing EDIII (rOMV-expressing EDIII from all four serotypes (rOMV(tEDIII)) with a dosage of 10 µg of expressed antigen in the OMVs and 10 µg of soluble purified antigens (tEDIII). The volume injected in each mouse was 100 µl. Fifteen days post-priming, blood samples were collected from the lateral tail vein of the mice. PBMCs and serum samples were isolated for profiling of T and B-cell responses. On day 15, the mice from all experimental groups were administered a booster dose identical to the primary immunization. On day 28 post-priming dose, mice were euthanized by asphyxiation through placement in a chamber and introduction of 100% CO2 at a fill rate of 30-70% displacement of the chamber volume per minute, added to the existing air in the chamber. Subsequently, blood, spleen, and lymph nodes were harvested for various immunological assays.

### Recall assay for antigen specific T-cell response by Flowcytometry

Antigen-specific T cell responses were analyzed by restimulation of splenocytes after 28th days after immunization. In brief, spleens from different groups of mice were isolated at 28th day post immunization, and a single suspension was prepared by mechanical disruption along with filtration with a 70µm strainer to obtain a mono cell suspension. Cells were pelleted at 360xg six min, and the obtained pellet was resuspended in 1X RBC lysis buffer (BD Biosciences) for 10 min at room temperature to lyse the red blood cells. A single suspension of cells was seeded in a bottom 96 well plate at a density of 0.5 x 106/well and treated with DENV-2 antigen (10µg/ml) for 10 h in the presence of Golgi Stop (BD Biosciences). After treatment, the cells were pelleted and washed twice with FACS buffer (BD Bioscience) and stained with anti-CD4-PerCP and anti-CD8-FITC (1:300) for 1 h at RT. Now, cells were washed and fixed with 4% PFA for 20min @ RT, followed by permeabilization using perm buffer (Biolegend USA) for another 20 min @ RT for intracellular staining. Antibodies against IFNg- and IL-2 (anti-IL2-PE and anti-IFNg-APC, respectively) were added (1:250) and incubated for 1 h at RT. Finally, cells were washed and resuspended in FACS buffer, data were acquired using flow cytometry (BD Accuri C6), and data were further analyzed in Flowjo (BD bioscience).

### Serum Antibody ELISA

96 well ELISA plate (maxisorp-NUNC) was coated with 100 μl/well of DENV EDIII protein (10 μg/ml) from all four serotypes in bicarbonate coating buffer (pH 9.5) and incubated at 40C overnight. Plates were washed three times in washing buffer (1XPBS + 0.05% tween-20) and blocked with 200 μl of 4% skimmed milk for 2 h at RT and further washed thrice. Serum samples were diluted in 0.1% skimmed milk prepared in 1× PBST, added to the wells, and incubated for 2 h at RT. After incubation, the plates were washed four times and incubated with an HRP-conjugated anti-mouse secondary antibody for 1 h at RT. Plates were washed five times and TMB substrate 50 μl/well were added in the plate, after certain time-points reaction was stopped using 0.2 N H2SO4, finally absorbance were taken at 450 nm and plotted on Y axis. In another set of experiments to evaluate the antibody titer, antigen-coated plates were incubated with various dilutions of pooled serum from 10 mice in two groups of five each for 2 h at RT; the remaining steps were the same as above.

### Preparation of DENV Stocks

DENV 1 (Hawaii), DENV 2 (TR1751), DENV 3 (Thailand 1973) and DENV 4 (Columbia 1982) viral strains were propagated in C636 cells, as described previously [20]. These cells were cultured in α-MEM media supplemented with 10% FBS and 1% antibiotic/antimycotic (Gibco) in T75 flasks. Cells were infected with the virus (0.1 moi) for 30 min at 33°C. The inoculum was removed, and fresh media with reduced FBS was added to the cells and kept for five–seven days until the cytopathic effect (CPE) was observed. The viruses were harvested by centrifugation, and the supernatants containing the viruses were stored at −80°C. Virus titers were quantified by FACS titration assay in Vero cells, as described elsewhere[21].

### Flowcytometry Based Neutralization Test (FNT)

Serum samples from immunized mice were assessed for neutralizing antibodies against DENV using the FNT method[21]. To evaluate the neutralizing capacity of sera, an in-vitro infection model was replicated in which Vero cells were cultured in DMEM media containing 10% FBS and 1% antibiotic/antimycotic (Gibco Life Technologies USA) and seeded in a 96-well plate at a density of 2.5×104 cells/well. These cells were then incubated with a mixture of heat-inactivated serum samples from 10 mice in two groups of five each and different dilutions of dengue virus from various serotypes. After 2 h, the virus-serum mixture was removed and 200 μl of DMEM media containing 2% FBS was added to each well. The plates were then incubated at 37°C with 5% CO2 for 24 h. Following incubation, the infected cells were washed with 1× PBS and stained with an anti-DENV (D1-4 GeneTex USA) antibody (1:500) for FACS analysis, as previously described. The stained samples were analyzed by flow cytometry by capturing 20,000 cells per sample. The percentage reduction in infection for each serum dilution was calculated, and the serum titers were represented as the reciprocal of the serum dilution that inhibited 50% of the virus (FNT50)[21, 22].

### Statistical Analysis

All data are presented as the mean ± S.E.M of three or four independent experiments. Unpaired Student’s t-test with Mann-Whitney post test was performed using GraphPad Prism software to evaluate the significance of differences between groups. Statistical significance was set at p < 0.05.

## Ethical statement

The experiments were executed according to protocols approved by the institutional biosafety committees of the University of Hyderabad.

## Data availability

This study did not include data from external repositories. All relevant data are included in the manuscript and supporting information files.

## Acknowledgments

The Authors would like to thank all the facilities provided by the School of Life Sciences, University of Hyderabad, including the flow cytometry and Transmission Electron Microscopy core facilities

## Author Contributions

**SD:** Investigation, designed and performed experiments, analyzed data, and edited the manuscript. **FA:** Manuscript writing (original draft) and editing, experiments performed, and data analysis. **RAK:** Conceptualization, performed experiments, and analyzed the data. **SMR:** Responsible for constructing the EDIII-ClyA fusion product. **SN:** Animals and flow cytometry-related experiments. **AMP:** Conducted flow cytometry-related experiments; **DP:** Assisted by vector construction and OMV purification. **AA:** Provided valuable input to the immunology section and critical reviews of the manuscript. **NK** Conceived the research idea and supervised and contributed to manuscript editing and funding acquisition. All authors have read and approved the final draft.

