## Supplementary Material for "Programing Immunogenicity of Dengue EDIII Vaccine Antigens Using Engineered Outer Membrane Vesicles (OMVs)"

**Supplementary Data**

**Figure S1**- Sub cloning of EDIII antigens into pBAD-ClyA-Hist vector to produce EDIII-ClyA fusion product.

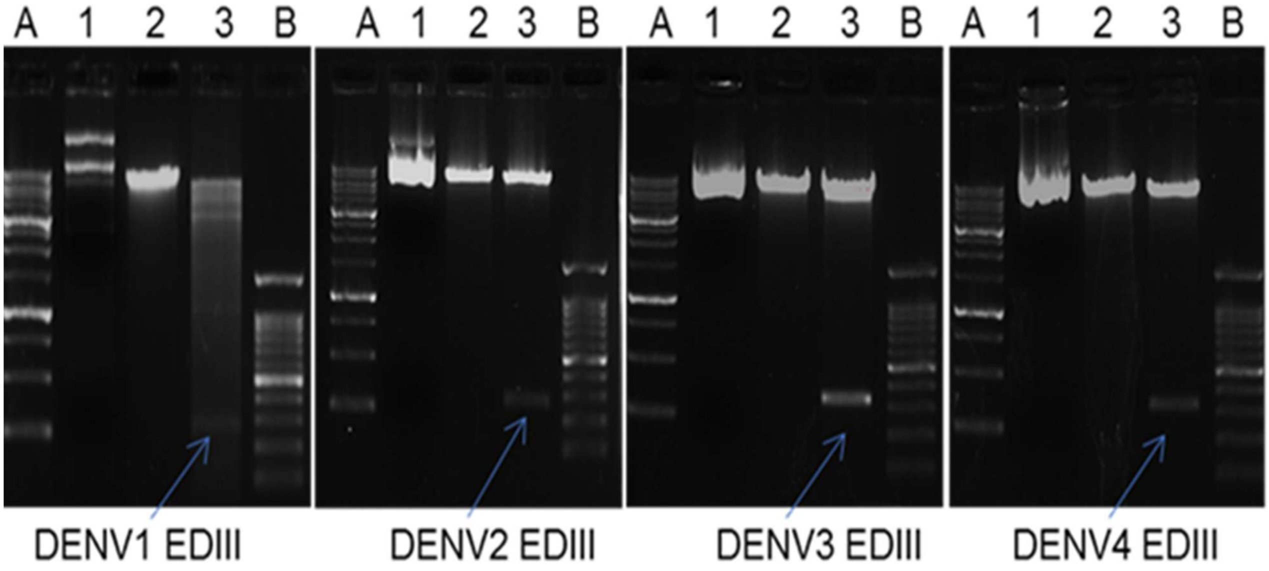


Figure S1. Data demonstrates the cloning of EDIII domain in ClyA-pBAD vector. The indicated EDIII from DENV1-4 were cloned into ClyA-pBAD vector. DNA from indicated clones were digested with XbaI and HindIII and separated by agarose gel electrophoresis. 1) Uncut; 2) Single cut with Hind III; 3) Double digested with XbaI and HindIII ; A) 1 Kb ladder; B) 100bp ladder

**Figure S2.** rOMVs were characterized for the presence of EDIII antigen fused with ClyA using western blotting in which blot were probed with anti-His and antigen-specific anti-sera.

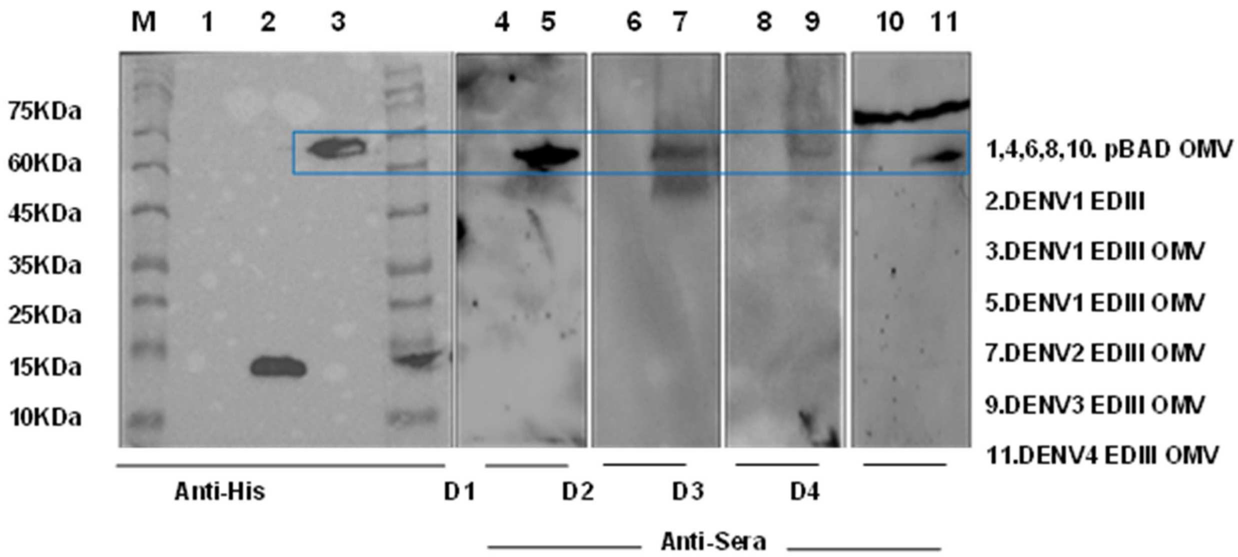


Figure S2. Data demonstrates the western blotting confirmation of DENV antigens by anti-sera from immunized mice M- protein marker 2- recombinant EDIII protein (Anti His) and 5,7,9,11 are DENV EDIII 1 to 4 respectively

**Figure S3**. Antigen-presenting cells expressing co-stimulatory markers upon treatment. Representative FACS plots for co-stimulatory molecules like CD80, CD86, MHC I and MHC II.

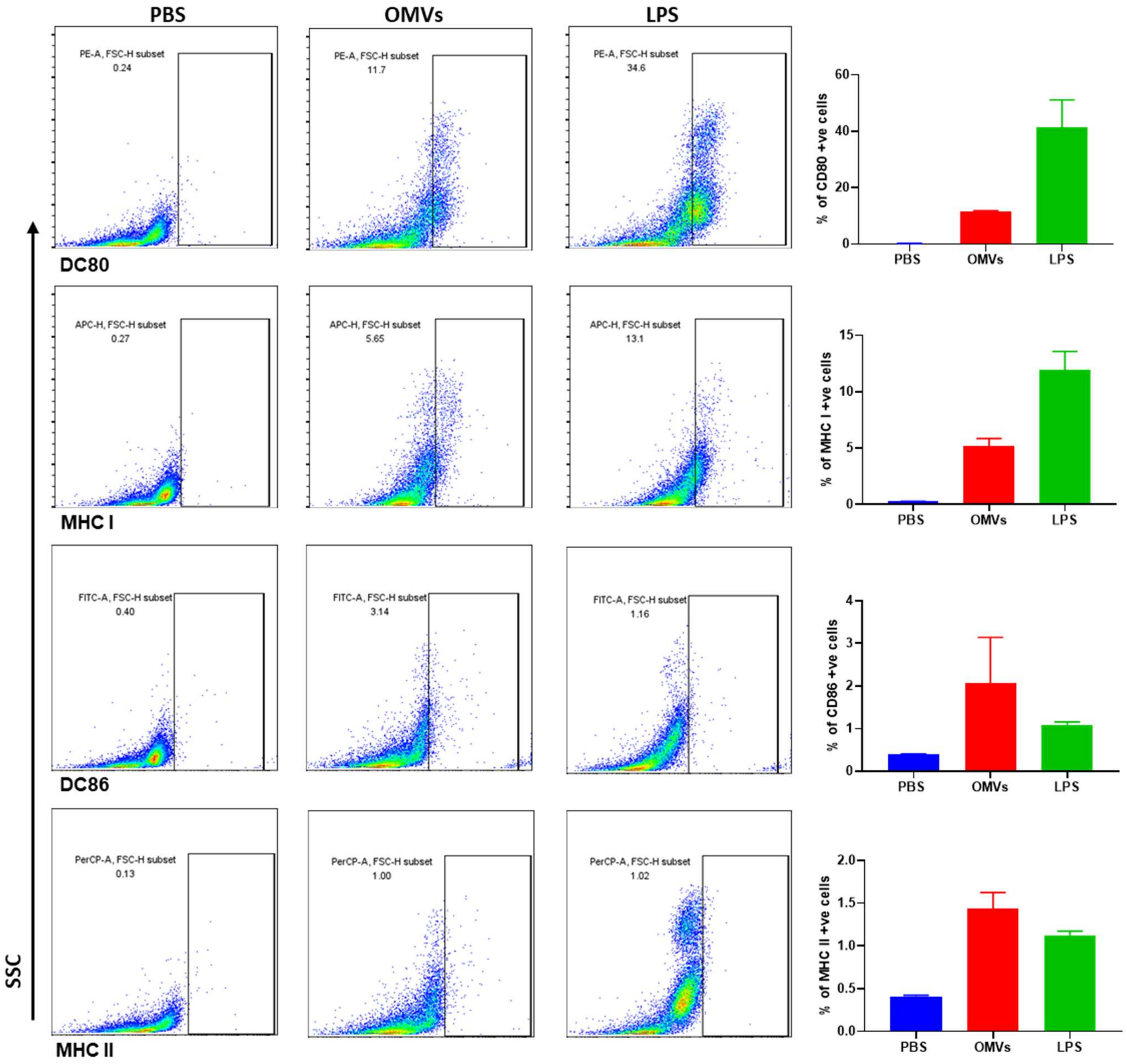

**Figure S4** Uptake of GFP-OMVs by antigen-presenting cells at $4^{0}$C and $37^{0}$C. Gating strategy and representative FACS plots of DCs with GFP-OMVs.

**Uptake at $4^{0}$C**

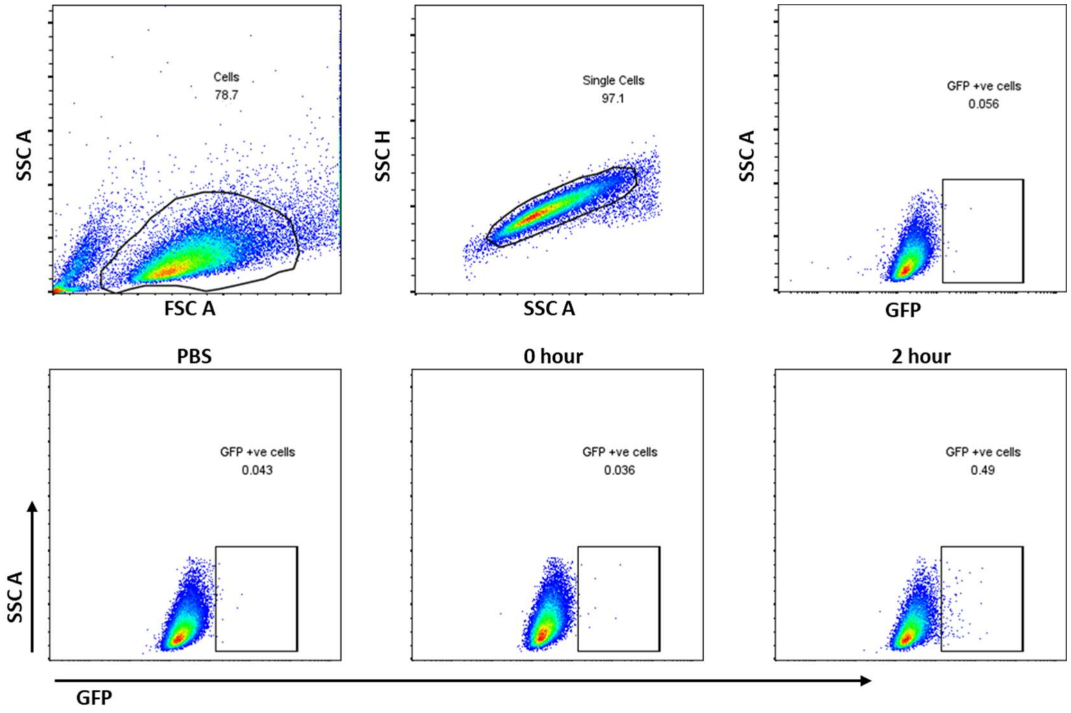


**Uptake at $37^{0}$C**

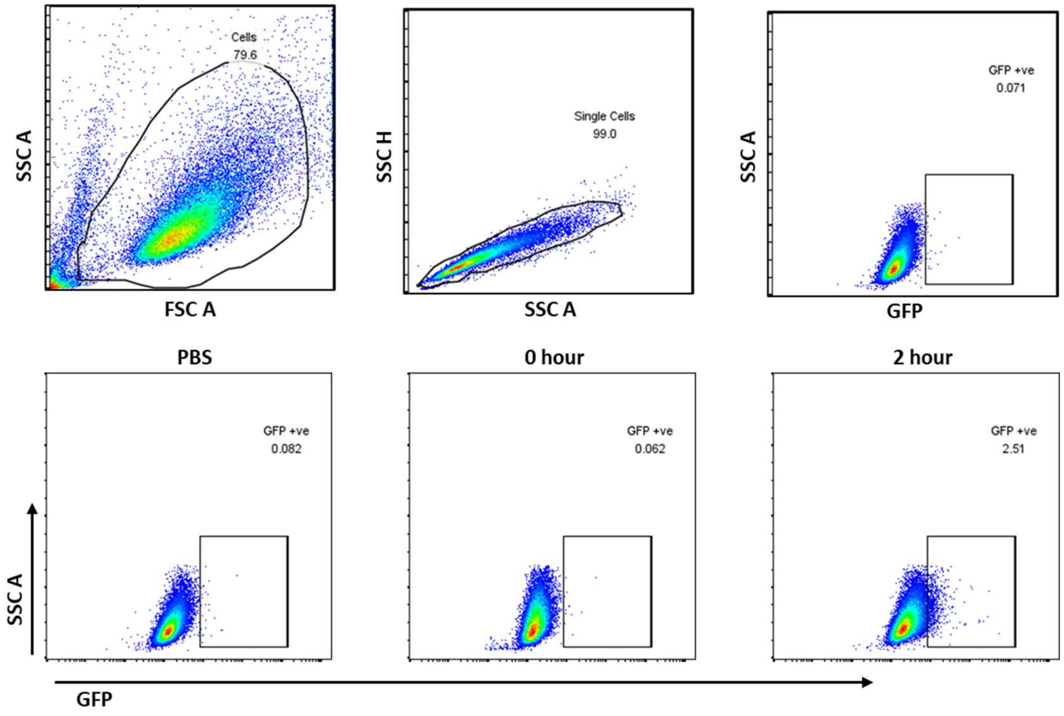

Figure S4. Antigen-presenting cells (DCs) were incubated with GFP-expressing rOMVs at two different temperatures (4°C and 37°C). The percentage of GFP-positive cells was evaluated using flow cytometry (FACS) with gating, demonstrating significant rOMV uptake at 37°C while no uptake was observed at 4°C.
